# Novelty-seeking and directed exploration scale with dissociable social network outcomes and brain systems in older adults

**DOI:** 10.64898/2026.09.21.753261

**Authors:** Christian Valtierra, Jeremy Hogeveen, Gabriella Mace, Samantha Moreno, Avery Ostrand, Sydney Griffith, Maria José Auil, Patrick McConnell, Joseph Chen, Jennifer Mitchell, Gary R. Turner, R. Nathan Spreng, Adam Gazzaley, Lorenzo Pasquini

## Abstract

Social isolation and loneliness are major determinants of well-being in older adults. Shrinking social networks and age-related shifts in social behavior and decision-making may increase vulnerability to loneliness. These changes in social decision-making strategies can be conceptualized within an exploration-exploitation tradeoff framework, in which social options with known outcomes are prioritized over uncertain ones. Yet, no study has characterized how complementary decision-making strategies, undirected novelty seeking and goal-directed exploration, relate to social network constructs or are supported by distinct neural systems. These strategies capture distinct forms of decision-making: undirected novelty seeking is driven by the intrinsic salience of novel options independent of expected informational gain, whereas goal-directed exploration involves deliberately sampling uncertain options to reduce uncertainty and optimize future choices. Here, we used fMRI and a novel social bandit reinforcement task to identify dissociable social constructs and brain systems linked to undirected novelty seeking and goal-directed exploration in older adults. Undirected novelty seeking was associated with social network size and with activity of temporoparietal and mid-cingulate brain areas. Goal-directed exploration predicted subjective feelings of social connectedness and was associated with activity of anterior cingulate and prefrontal brain areas. Crucially, choice probabilities from a non-social version of the three-arm bandit task were not associated with social decision-making nor with social constructs. Our findings suggest that, in later life, dissociable social decision-making strategies, when faced with unknown options, rely on distinct neural systems supporting social network size as well as subjective feelings of social connectedness.

**Significance statement:** Loneliness and social isolation are major threats to healthy aging. Understanding social decision-making strategies and their neural underpinning in older adults is essential to develop novel interventions targeting these major risk factors for negative outcomes in later life. Using fMRI and a social reinforcement bandit task, we show that undirected novelty seeking is linked to social network size, while goal-directed exploration is linked to subjective feelings of social connectedness. Both behaviors rely on distinct, distributed brain networks spanning brain areas relevant to decision making and social cognition.

## I. Introduction

Social relationships are a central determinant of emotional well-being and health in older adulthood. Across the lifespan, social networks undergo systematic restructuring, with older adults maintaining fewer but emotionally closer relationships (Bruine De Bruin et al., 2019; Carstensen, 1992; Rook & Charles, 2017). This transition can be conceptualized as a shift in social decision-making strategies, where the informational value of new relationships declines (Spreng & Turner, 2021) and the emotional value of closeness with familiar others increases (Carstensen et al., 1999). This shift, although supporting emotional regulation and well-being under stable circumstances, mayClick or tap here to enter text. increase vulnerability to social isolation when social networks are disrupted by late-life events such as retirement, relocation, bereavement, or declining health (Anusic & Lucas, 2013; Freak-Poli et al., 2022). Reduced social engagement is strongly associated with loneliness, depression, cognitive decline, and increased mortality risk. This may create a self-reinforcing cycle that undermines well-being and resilience and increases vulnerability to neuropsychiatric and neurological disorders (Cacioppo & Cacioppo, 2014; Hawkley & Cacioppo, 2010). To prevent this cascade, it is essential to identify the behavioral and neural mechanisms that support adaptive social engagement and promote prevention-oriented approaches for healthy aging.

The tendency of older adults to prioritize relationships with familiar others over the formation of new social connections can be conceptualized as an exploration– exploitation dilemma (Spreng & Turner, 2021), in which exploitation involves selecting the best option based on current knowledge, whereas exploration involves sampling new options that may lead to better future outcomes at the expense of an immediate exploitation opportunity (Daw et al., 2006; Hogeveen et al., 2022). Within this framework, two distinct strategies govern decision-making in the face of uncertain and novel options (Gershman, 2018; Wilson et al., 2014). *Undirected novelty seeking* (Blanchard & Gershman, 2018) reflects the bias to sample a previously unencountered option, independent of any explicit estimate of its uncertainty or expected value(Wittmann et al., 2008). In a complementary but separate strategy, *goal-directed exploration* balances information seeking and reward maximization (Cohen et al., 2007), where options with imprecise value estimates are prioritized to reduce uncertainty and optimize future choice. While undirected novelty seeking and goal-directed exploration are often correlated (because novel options are inherently uncertain) these two strategies reflect computationally and behaviorally dissociable processes relying on distinct neural systems (Cockburn et al., 2022). For example, dopaminergic midbrain– striatal circuits and parietal regions, including the temporoparietal junction (Corbetta et al., 2008; Corbetta & Shulman, 2002), are involved in salience detection and orienting toward novel stimuli, while dorsal anterior cingulate cortex and related noradrenergic systems track uncertainty and support information-seeking behavior (Aston-Jones & Cohen, 2005; Jepma & Nieuwenhuis, 2011; Kolling et al., 2012; Shenhav et al., 2014).

A large body of previous work has focused on decision-making in an abstract or non-social context (for a review, see Wyatt et al., 2024), limiting its ability to explain shifts in social behavior in aging. Here, we recruited a cohort of older adults and adapted a validated bandit reinforcement task to examine whether novelty seeking and exploratory behavior predict distinct facets of social network constructs and are supported by separable neural systems. We designed a multi-arm bandit reinforcement learning task using human faces as intrinsic social stimuli (Haxby et al., 2000) applied a partially observable Markov decision process model (POMDP) (Averbeck, 2015; Hogeveen et al., 2022) to derive measures of social goal-directed exploration and undirected novelty seeking. We linked these measures to psychometric assessments and functional imaging data, revealing that undirected novelty seeking predicts social network size and is associated with activity of posterior brain areas, while goal-directed exploration predicts social network quality and relies on activity of frontal brain areas.

## II. Methods

### II.a. Study sample

The study protocol was approved by the University of California, San Francisco (UCSF) Institutional Review Board (22-37461) and was conducted in accordance with the Declaration of Helsinki. Informed consent was collected for all participants. The study was registered as part of a larger clinical trial (https://clinicaltrials.gov/study/NCT05645835).

Twenty-five cognitively healthy older adults (ages 60 – 85 years) (**Table 1**) were enrolled from a database of older adult control volunteers, who had been assessed with neuropsychological tests of executive and memory function via a previously described remote characterization module (Arioli et al., 2022) at the UCSF Neuroscape Center. Recruitment and study design aimed to reduce between-subject heterogeneity and measurement error, thereby maximizing the expected effect size of individual-differences associations (DeYoung et al., 2025). Specifically, we recruited a well-characterized, cognitively healthy sample and used standardized behavioral and neuroimaging measures. The neuropsychological evaluation administered working memory and verbal learning tests, including the CVLT-II (Delis et al, 1987), processing speed (WAIS-R) (Wechsler, 1997), visual-motor sequencing (DKEFS Trail-Making A and B), semantic fluency and phonemic fluency (DKEFS) (Delis et al., 2001). Participants had to score within one standard deviation of the age-adjusted mean to be considered cognitively healthy and included in the study. English fluency and normal or corrected-to-normal vision were required to be included in the study. Important exclusion criteria included MRI-contraindications and the self-reported presence of major neuropsychiatric, neurological, or systemic diseases. Recorded demographic variable included age, assigned sex at birth, years of education, and household income.

**Table 1.** Participant characteristics and demographics. Demographic characteristics of the 25 participants enrolled in the study.

Figures, Tables, and Legends
| Participant Demographics (n=25) |  |
| --- | --- |
| Age in years | 70.8 ± 6.9 |
| Sex | 14 M, 11 F |
| Education in years | 18.2 ± 2.0 |
| Income in US \$ | |
| Below \$100,000 | 4 |
| \$100,000 - \$200,000 | 15 |
| \$200,000 - \$300,000 | 3 |
| \$300,000 - \$400,000 | 2 |
| Above \$400,000 | 1 |
| Race and Ethnicity |  |
| White/Hispanic/Asian | 18/1/6 |
| Neuropsychological Surveys (mean ± SD) |  |
| UCLA Loneliness 20 (n=24) | 7.8 ± 7.1 |
| Social Connectedness Scale (n=24) | 45.0 ± 5.6 |
| Lubben Social Network Scale (n=24) | 33.8 ± 8.2 |
| Social Network Index (n=24) | 19.8 ± 12.8 |
| Remote Neurological Assessment (z) |  |
| CVLT Immediate Recall | 0.7 ± 1.0 |
| CVLT Short | 1.2 ± 0.8 |
| Digit Span Forward | -0.4 ± 0.9 |
| Digit Span Backward | -0.2 ± 0.7 |
| Lexical Fluency | 0.0 ± 0.9 |
| Semantic Fluency | -0.4 ± 0.9 |
| Trails | 0.0 ± 1.1 |
| CVLT Long | 1.0 ± 0.9 |
| CVLT Cued | 0.7 ± 1.5 |
| fMRI Head Motion Parameters (mean ± SD) |  |
| Framewise Displacement (mm) | 0.17 ± 0.6 |

### II.b. Social Constructs

Participants completed a battery of validated self-report questionnaires characterizing subjective and structural dimensions of social functioning. Subjective loneliness was assessed using the 20-item UCLA Loneliness Scale (Version 3), in which respondents rate feelings of disconnection from others on a 4-point Likert scale, with higher summed scores indicating greater loneliness (Russell, 1996). Subjective social connectedness was indexed using the Social Connectedness Scale (SCS), which measures the degree to which an individual feels connected to others in their broader social environment, with higher scores reflecting stronger belongingness (Lee & Robbins, 1995). Structural social integration was quantified using the 12-item Social Network Index (SNI), which captures participation across 12 types of social relationships (e.g., spouse, family, friends, neighbors, workmates, religious and community groups), yielding indices of network diversity and size, as well as a total score, based on whom participants have remote or in-person contact with at least once every two weeks (Cohen et al., 1997). Finally, given the specific relevance of family and friendship ties to social isolation in late life, participants completed the abbreviated 6-item Lubben Social Network Scale (LSN), a validated screening instrument developed for older adults that assesses the size, closeness, and frequency of contact within kin and non-kin networks, with lower scores indicating greater risk for social isolation (Lubben et al., 2006).

### II.c. Social-decision making task

After informed consent was obtained, participants were invited to an in-person fMRI visit during which they performed a three-arm bandit social decision-making task while their Blood-Oxygenation-Level-Dependent (BOLD) activity was recorded in the scanner. For this experiment, we adapted a previously validated three-armed bandit task which was successfully used to investigate exploration–exploitation tradeoffs in a non-social decision-making study involving both human subjects and macaques (Hogeveen et al., 2022).

Participants made speeded (≤2 seconds) manual responses among three pictures of faces of older adults taken from the Lifespan Database of Adult Facial Stimuli (Minear & Park, 2004) (https://agingmind.utdallas.edu/). Human faces were selected as stimuli because they represent socially salient stimuli that rapidly convey identity, emotional, and interpersonal information (Haxby et al., 2000, 2002; Todorov, 2008). These faces have been rated in previous work for levels of likeability and trustworthiness by two separate research groups (Ebner & Ebner, 2008; Ramos et al., 2016). We selected 150 frontally oriented faces of older adults aged ≥55y, ensuring a gender balance between male and female faces (50%), and racial distributions aligned with the demographic characteristic of the general US population. Only faces showing neutral expression, based on the published descriptions of the faces, were selected. All pictures were visually checked for artifacts by three study team members (MJA, CV, and GM).

Stimuli were presented using PsychoPy v2023.2.2 (https://www.psychopy.org/), with responses recorded using a four-button MRI-compatible box (Current Designs, PA). Faces were randomly assigned an *a priori* low (*p*=0.2), medium (*p*=0.5), or high (*p*=0.8) reward probability. Reward probabilities were randomly assigned to faces for each participant. Every 5–12 trials (mean of 6 trials) a novel insertion took place, wherein one familiar face in the current set was replaced by one novel face to create a new set that would be presented for the successive 5–12 trials. Novel faces were randomly assigned a low, medium, or high reward probability, with the caveat that at least one out of the three face pictures could not have the same assigned reward probability as the other two face pictures in the new set. Participants completed 180 trials containing 23 novel stimulus insertions, divided evenly into three ∼9-minute fMRI runs. Face picture location was randomized on each trial, and participants received either reward (green ‘+1’) or non-reward (red ‘0’) feedback after each decision. Notably, jittered inter-stimulus intervals (both between face pictures and reward cues and between reward cues and face selection trials) were used to introduce random, variable timing between stimuli to enable the separation of overlapping hemodynamic BOLD responses, enhancing the ability to distinguish neural activity between closely spaced trials (Dale, 1999).

### II.d. Behavioral data analysis: empirical and modelled data

Choices made within two trials from an insertion were used to define best, worst, or undirected novel choices. Following an approach validated in previous studies (Hogeveen et al., 2022), the first two trials following an insertion were selected to maximize sensitivity to undirected novelty-driven behavior before reward contingencies could be reliably learned from experience. Best and worst choices were defined based on their inherent reward probabilities, while undirected novel choices were based on whether the new face picture was selected, regardless of their reward probabilities.

In addition, a partially observable Markov decision process (POMDP) formalizing decision-making under uncertainty was applied to empirical choice and reward data across all trials to model goal-directed explorative and exploitative choices. In a POMDP, individuals are assumed to make choices based on latent states of the environment that cannot be directly observed but must be inferred from accumulated choice and reward history. The model maintains a belief state, a running estimate of the reward probability and sampling count for each option, which is updated after every choice and outcome. From this belief state, the POMDP derives two values for each available option on each trial: (i) the immediate expected value (IEV), reflecting the learned reward probability of a given option based on prior experience, and (ii) the future expected value (FEV), reflecting the discounted future gains expected from the latent choice sequences that could follow. IEV therefore indexes the value of exploiting a known option. To isolate the value of exploring, we computed an exploration BONUS for each option, defined as the difference between that option’s FEV and the average FEV across all available options in the current set. To estimate the relative weighting of these value terms on choice, the model-derived IEV and BONUS estimates for all options were passed through a softmax choice function with two participant-specific free parameters, one scaling IEV and one scaling BONUS, fitted separately for each individual to yield trial-by-trial choice probabilities across the choice set. Explorative and exploitative trials were then classified directly from the POMDP-derived estimates of the chosen option: explorative trials were defined as those where IEV < 0.5 and BONUS > 0, reflecting a choice driven more by information-seeking than by known reward value, and exploitative trials were defined as those where IEV > 0.5 and BONUS < 0, reflecting a choice driven by known reward value at the cost of future information gain. These criteria were selected to identify trials in which choice behavior was predominantly driven by information-seeking (goal-directed exploration) versus known reward value (exploitation), and trials meeting neither criterion were classified as neither explorative nor exploitative. Reward prediction errors (RPEs) were calculated by subtracting the IEV from the observed binary reward outcome (coded as 1 for reward and 0 for non-reward), yielding positive prediction errors when outcomes exceeded expectations and negative prediction errors when outcomes fell below expectations.

One-way ANOVAs (*p* < 0.05) and post-hoc paired comparisons (uncorrected and FWE-corrected *p* < 0.05) were used to compare mean reaction times and individual choice probabilities across the distinct choices. Individual ratios of best and novel choices were calculated by dividing the probability that a participant chose a novel face picture by the overall probability of choosing the picture with the highest reward of probability. Similarly, individual ratios of goal-directed exploration and exploitation were calculated by dividing the probability that a participant explored by the overall probability of exploiting. Ratios were used to capture the relative balance between competing decision-making strategies while accounting for individual differences in overall choice tendencies. Partial Pearson’s correlations corrected for age, sex, and years of education were used to investigate the relationship between choice ratios and social network constructs (*p* < 0.05).

### II.e. Remote animal-decision making task

A subset of participants (n = 16) remotely completed a web-based follow-up task administered through lab.js on their personal laptops (Henninger et al., 2021). The task was designed to isolate the contribution of social information to explore–exploit behavior by pairing a structurally matched non-social bandit against the face-based paradigm; whereas the social bandit task used face stimuli, which are inherently social, the follow-up task used mages of animals. Specifically, stimuli were a subsample of the “animal” concept category drawn from the THINGS database of naturalistic object images (Hebart et al., 2019), for a total of 110 images.

The task, consisting of four blocks, each approximatively 5 min in length, was adapted from a previously validated three-armed novelty bandit (Hogeveen et al., 2022) and retained its overall structure: 224 trials divided into 4 blocks of 56 trials each. On each trial, participants viewed three animal images for a maximum of 2 seconds and selected one. Unlike the MRI task, stimulus presentation transitioned directly into the inter-stimulus interval upon response. Following the inter-stimulus interval, probabilistic binary feedback (reward or no reward; “+1” or “0”) was delivered for 1 sec according to fixed per-option reward probabilities of 0.2, 0.5, and 0.8. Analogous to the social version of the bandit-task, at pseudo-random intervals, a novel image was introduced in place of one existing option, allowing exploration of newly available stimuli to be dissociated from exploitation of familiar ones. The four blocks were independent from each other: at each block boundary, all three options were replaced with a new set of images, so no stimulus or value information carried across blocks. A start-of-trial fixation cross (serving as the inter-trial interval) and the post-choice inter-stimulus interval were each jittered uniformly between 500 and 2000 ms (grand mean inter-trial interval = 1.2 seconds; grand mean inter-stimulus interval = 1.2 seconds).

The behavioral data generated through this non-social version of the three-arm bandit task was analyzed analogously as the social version using faces of older adults. Probabilities for best and novel choices and for goal-directed exploration and exploitation were derived from each task block and averaged, before computing ratios of novel/best and exploration/exploitation for each individual. Partial Pearson’s correlations corrected for age, sex, and years of education were used to investigate the relationship between non-social choice ratios and social choice rations, as well as between non-social choice ration and social network constructs (*p* < 0.05). Differences in partial correlations between social and non-social choice rations with psychometric measures were assessed using Williams’ test for dependent overlapping correlations (*p* < 0.05), accounting for the correlation between social and non-social choice ratios within the same participants

### II.f. Neuroimaging data acquisition

Participants were placed in a 3T Siemens Prisma Fit (Siemens, Germany) at the UCSF Neuroscience Imaging Center with a 64-channel head coil. Noise cancelling headphones and earplugs were used to mitigate noise from the scanner. A four-button box was used to choose between face pictures during the social bandit task. The scan session started with a structural MPRAGE scan (TR = 2,300 ms, TE = 2.96 ms, TI = 900 ms, flip angle = 9°, isotropic voxel sizes of 1.0 mm) of approximately six minutes. This was followed by the participant performing a practice version of the social bandit task. During each of three social bandit task blocks, multi-echo, multiband T2*-weighted BOLD EPI data were acquired using three echoes, a multiband acceleration factor of 4, and an in-plane GRAPPA acceleration factor of 2. Each run comprised 554 time points (554 volumes per echo; acquisition time = 9 min 30 s), with 48 axial slices acquired in interleaved order, anterior-to-posterior phase encoding, 2.6-mm isotropic voxels, TR = 1,000 ms, and TEs = 13.40, 32.12, and 50.84 ms. The task was synchronized to the scanner and fieldmaps were acquired to improve distortion correction and improve registration of fMRI data.

### II.g. Neuroimaging data preprocessing

Functional MRI data were preprocessed using fMRIPrep v23.2.0a3 (Esteban et al., 2019). Structural images underwent bias-field correction, skull stripping, tissue segmentation, and cortical surface reconstruction using ANTs and FreeSurfer. Nonlinear spatial normalization to MNI152NLin6Asym space was estimated using ANTs. Functional images were corrected for head motion (MCFLIRT, FSL), coregistered to the T1-weighted anatomical image using boundary-based registration (bbregister, FreeSurfer), and resampled using a single interpolation step combining all estimated transforms. The normalization transformation matrices were stored for future use. Nuisance regressors, including framewise displacement, white matter, and CSF signal estimates, were extracted for use in subsequent denoising analyses. Motion outliers were identified using thresholds of mean framewise displacement > 0.25 mm.

We subsequently used Tedana (TE-Dependent ANAlysis) (Dupre et al., 2021), an open-source Python library designed for multi-echo fMRI processing (https://tedana.readthedocs.io/en/stable/). Tedana leverages differences in signal decay across echo times to optimally combine multi-echo acquisitions, increasing BOLD sensitivity and signal-to-noise ratio. Optimally combined multi-echo BOLD timeseries in native space were generated and subsequently smoothed (6 mm FWHM kernel) and high-pass filtered using discrete cosine drifts with a 128 sec cutoff in Nipype (https://nipype.readthedocs.io/en/latest/). The resulting optimally combined BOLD timeseries were then used for first-level analyses in native space.

### II.h. First- and second-level models

First-level analyses were conducted in native space using three distinct general linear models (GLMs) implemented in Nilearn v0.11.1 (Abraham et al., 2014; https://nilearn.github.io). The three models were constructed to serve complementary analytic goals: a quality-control model to verify that the pipeline recovered canonical stimulus-evoked responses; an event-related model to capture trial-wise behavioral events surrounding novel-option insertions; and a parametric model to test the neural correlates of latent decision variables derived from the POMDP model. Each regressor was convolved with the canonical SPM hemodynamic response function (Friston et al., 1998) as implemented in Nilearn. In the quality-control model, task events were modeled as boxcars matching their true presentation durations (see below); in the event-related and parametric models, all events were modeled as zero-duration stick functions time-locked to event onset.

In addition to the model-specific regressors of interest, each GLM included the following nuisance regressors: eight cosine drift regressors, implementing a high-pass filter with a cutoff of 1/128 seconds, six head-motion parameters (three translations and three rotations), and framewise displacement, for a total of sixteen regressors including the intercept. The resulting subject-level beta maps were then resampled from native space into standard MNI152NLin6Asym 2-mm space using ANTs with Lanczos interpolation, and the subject-specific spatial-normalization transforms previously estimated by fMRIPrep. The three first-level models were specified as follows.

#### Quality-control model

Task events were modeled as boxcars: *(i)* the fixation cross (jittered duration), *(ii)* face stimuli (2 s), *(iii)* rewarded feedback (1 s), and *(iv)* non-rewarded feedback (1 s).

#### Event-related model (early choices at insertion)

All events were modeled as stick functions: *(i)* regressors for best, worst, and novel choices made within two trials of an insertion; *(ii)* regressors for rewards and missed rewards at insertions; and *(iii)* a regressor for choices made more than two trials after an insertion, together with a regressor for the associated reward cues.

#### Parametric model

On each trial, the chosen option’s IEV and BONUS terms, derived from the POMDP model, were used to classify the choice as exploratory, exploitative, or neither. A choice was classified as exploratory when its BONUS term was positive and its IEV term was at or below 0.5 (BONUS > 0 and IEV ≤ 0.5), and as exploitative when its IEV term exceeded 0.5 and its BONUS term was negative (IEV > 0.5 and BONUS < 0); all remaining choices were assigned to the “neither” category. Exploratory and exploitative choice onsets were each modeled as stick-function regressors, the “neither” regressor was included as a nuisance regressor of no interest. Reward-prediction-error events were modeled as separate stick-function onset regressors carrying parametric modulators for positive (PRPE) and negative (NRPE) prediction errors. Finally, a choice-locked novelty regressor carried the number of trials elapsed since the most recent novel insertion, mean-centered within run, such that a negative slope indexes a novelty-responsive response that declines with increasing familiarity.

#### Second-level analyses

Six second-level random-effects analyses were estimated in SPM12 (Wellcome Centre for Human Neuroimaging; http://www.fil.ion.ucl.ac.uk/spm/), by entering beta maps from the first-level models averaged across the three runs into paired t-tests, resulting in subject-level contrast images. From the QC model we tested (1) face stimuli > fixation cross and (2) reward > no reward. From the event-related model we tested (3) best > novel choices and (4) rewards > missed rewards. From the parametric model we tested (5) exploitation > exploration and (6) positive > negative reward prediction errors. All analyses included age, sex, and education as co-variates of no-interest. For all models, voxel-wise statistics were thresholded at a cluster-forming threshold of p < 0.005 uncorrected, and inference was performed at the cluster level using Gaussian random-field theory with false-discovery-rate correction (FDRc) at p < 0.05. The Harvard-Oxford Cortical and Subcortical Atlas was used to describe the anatomical location of identified clusters (Desikan et al., 2006; Frazier et al., 2005; Goldstein et al., 2007; Makris et al., 2006).

### II.i. Neurosynth analyses

To characterize the cognitive-network distribution of neural activity associated with exploration and novelty, we performed a region-of-interest analysis using independently defined meta-analytic maps obtained from Neurosynth (Yarkoni et al., 2011). Association-test *z*-statistic maps thresholded at FDR-corrected *p* < 0.01 were extracted for the following terms: *(i)* social (1,302 studies, https://neurosynth.org/analyses/terms/social/), *(ii)* decision making (509 studies, https://neurosynth.org/analyses/terms/decision%20making/), and *(iii)* working memory (1,091 studies, https://neurosynth.org/analyses/terms/working%20memory/). These maps were binarized to define three independent network masks. For each participant, condition-specific beta estimates were used to generate subject-level difference maps for exploration versus exploitation (exploration − exploitation) and novel versus best choices (novel − best). Because the aim of this analysis was to quantify the magnitude of neural differentiation between behavioral conditions independently of its direction, the absolute value of the voxelwise beta difference was calculated. For each contrast and participant, network differentiation was quantified as the mean absolute beta difference across all voxels within each binarized Neurosynth mask. Thus, larger values indicate greater differentiation between the corresponding behavioral conditions within a given cognitive network, irrespective of whether individual voxels showed greater activity for one or the other condition. Repeated-measures ANOVAs (*p* < 0.05) and post-hoc paired comparisons with Holm correction (*p* < 0.05) were used to assess mean absolute beta differences across the decision making, social, and decision-making networks relative to working memory for both the exploration versus exploitation and novel versus best choices comparisons (repeated-measures *p* < 0.05 Holm-corrected; **Figure 4D-E**).

## III. Results

### III.a. Sample characteristics

Twenty-five cognitively healthy older adults (ages 60 – 85 years, 11 females) (**Table 1**) were enrolled in the study. All cognitive tests were within one standard deviation of the age-adjusted mean, suggesting intact cognitive functions. Mean framewise head displacement (FD) was used to assess the quality of fMRI data (Power et al., 2011), with all participants having a score below 0.25 mm, indicating low overall movement. All participants reported being right-handed. Psychometric social network constructs for one participant were lost due to a technical issue.

### III.b. Novel face selection correlates with social network size

The choices made by participants at each trial coupled with the known probability of reward for each face were used to identify empirical novel, best, and worst choices. Plotting participant-level averaged novel and familiar choices over time revealed that novel options were selected less often as the number of trials elapsed since a novel option was introduced, increasing instead the selection of the best available option (**Figure 2A**). Across the whole task, the participants’ overall probabilities for selecting novel, best, and worst options were calculated for the first two trials after insertion of a novel option, revealing that novel options were chosen more frequently than best options, with worst options being the less frequently selected choices (**Figure 2B**, F(2,48) = 16.98, *p* < 0.0005, post-hoc *t*-test *p* < 0.05 FWE corrected and uncorrected). Reaction times also differed across choices, with best options having the shortest reaction times when compared with novel and worst options, as shown in previous decision-making studies using a reinforcement learning bandit task (**Figure 2C**, F(2,48) = 4.01, *p* = 0.025, post-hoc *t*-test *p* < 0.05 FWE corrected and *p* < 0.1) (Hogeveen et al., 2022).

**Figure 1.**
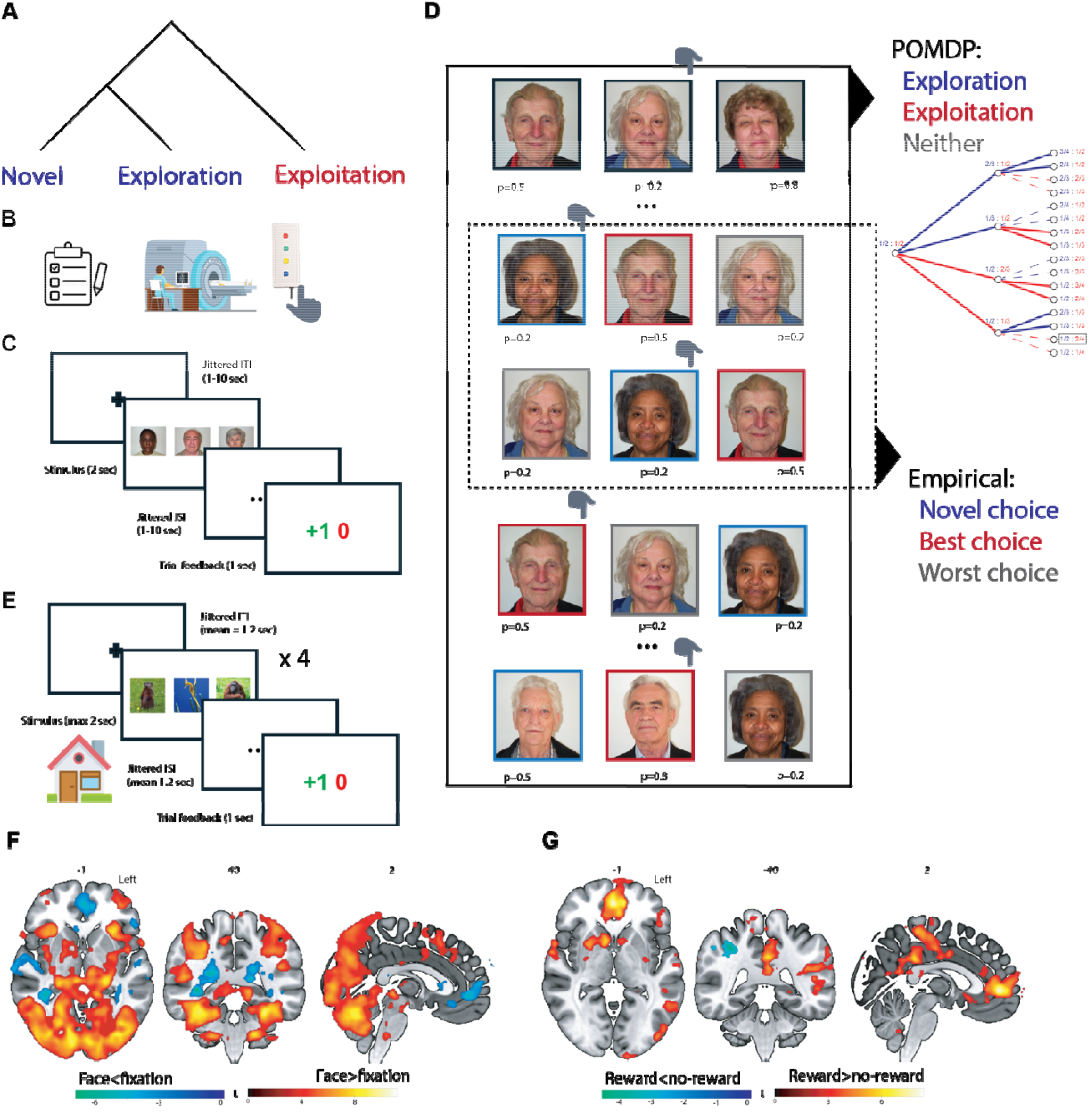
Study design and procedures. **(A)** When choosing between a familiar and a novel option, individuals may rely on different decision-making strategies. Exploitation refers to selecting options with known reward values. When options with uncertain outcomes are considered, two complementary strategies can be engaged. Undirected novelty seeking refers to the stochastic sampling of previously unencountered options, independent of their expected value, whereas goal-directed exploration reflects choices that balance information seeking with reward maximization. **(B)** After completing study surveys, participants were invited for an in-person fMRI visit, during which they completed a social multi-arm bandit task while their BOLD activity was measured in the scanner. **(C)** The multi-arm bandit reinforcement learning task consisted of 180 trials during which participants made speeded (2 sec) choices between three pictures of older adults. The inter-trial and inter-stimuli intervals (ITI and ISI) were jittered to optimize the deconvolution with the hemodynamic response, and short 1 sec reward cues were presented after each choice, indexed by a green +1 or a red 0. **(D)** Participants chose between pictures of older adults assigned a high, medium, or low probability of reward. Every 5-12 trial, a new picture was inserted, with all participants experiencing a total of 23 insertions. The two trials after a novel insertion (dashed box, mid right) were used to empirically measure novel (in blue), best (in red), and worst (in gray) choices. Choices made during the whole trial, as well as associated rewards, were implemented into a POMDP (undashed box, top right) to compute modelled choices for exploration (in blue), exploitation (in red), and neither (in gray). The dendrogram depicts a schematic representation of bandit options modelled by the POMDP. Blue lines (and fractions) indicate explorative choices, red lines (and fractions) indicate exploitative choices. The numerator and denominator of the fractions define the posterior probability of a reward. Thick lines show optimal choices; thin dashed lines show non-optimal choices [adapted with permissions from Averbeck 2015]. **(E)** After completing the MRI visit, participants completed a version of the multi-arm bandit task using animal pictures instead of faces of older adults as stimuli. Participants completed this version of the task remotely online. **(F)** The face versus fixation contrast, estimated across all trials, revealed activations within visual cortex, the fusiform cortex, the insula, the hippocampus, the temporoparietal junction, and the cerebellum (hot colors), while deactivations were notable in anterior cingulate cortex and auditory cortex (cold colors). **(G)** The reward versus no-reward contrasts, estimated across all trials, revealed activations within the medial prefrontal cortex, the ventral striatum, and medial parietal areas (hot colors), while a small cluster of deactivations was identified in the right parietal cortex (cold colors). Significant clusters were identified with a height threshold *p* < 0.005 and a cluster-level FDR-corrected threshold of *p* < 0.05; color bars show *t*-values. BOLD = blood oxygenation dependent signal; POMDP = partially observable Markov decision process

**Figure 2.**
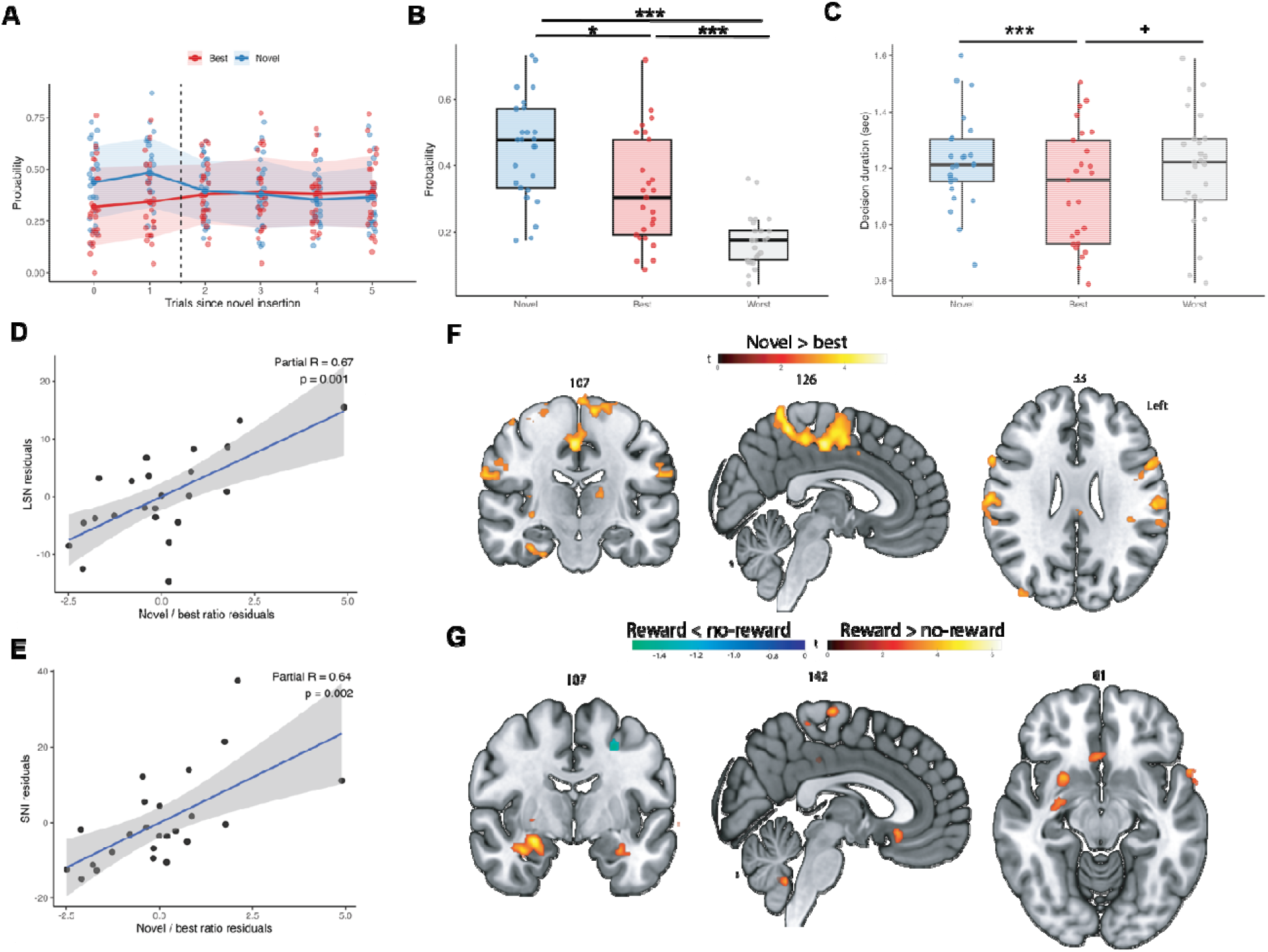
Face novelty seeking predicts social network size and is associated with activity in in mid-cingulate, supramarginal, and supplementary motor areas. **(A)** The novel option was selected less often as the number of trials elapsed since it was introduced, increasing instead the selection of the best familiar option. (B) The probabilities of choosing the novel, best, and worst option were computed for the first two trials after a novel insertion. Novel options were selected more often than best options, with worst options being selected the least frequently. (C) The individual-averaged reaction times for best options were significantly lower than for novel options and trending lower than for worst options. (D-E) The ratio of novel versus best options correlates positively with two measures of social network size, the Lubben Social Network (LSN) index and the Social Network Index (SNI). Partial Pearson’s correlation coefficients corrected for age, gender, and years of education. (F) Increased activity in the mid-cingulate cortex, supramarginal gyrus, and supplementary motor areas (warm colors) for novel versus familiar choices. (G) The reward versus no-reward contrasts for novel insertions revealed activations within the bilateral hippocampi, the basal forebrain, and the right ventral putamen (hot colors), while a small cluster of deactivations was identified in the left superior frontal cortex (cold colors). Significant clusters were identified with a height threshold *p* < 0.005 and a cluster-level FDR-corrected threshold of *p* < 0.05; color bars show *t*-values. ^+^*p* < 0.1; \**p* < 0.05; ***FWE-corrected *p* < 0.05

We next assessed whether choice selection was associated with perceived properties of the selected faces. To address this question, we leveraged out-of-sample scores of likeability, attractiveness, and trustworthiness published by two distinct studies (Ebner & Ebner, 2008; Ramos et al., 2016). These scores were correlated with the faces’ probabilities of being selected, calculated over all trials and averaged across study participants. The average selection probability of a face did not significantly correlate with scores of attractiveness (R_Ramos_(54) = 0.20, *p* = 0.147; R_Ebner_(48) = 0.09, *p* = 0.538), likeability (R_Ramos_(54) = 0.16, *p* = 0.224; R_Ebner_(48) = −0.05, *p* = 0.710), or trustworthiness (R_Ramos_(54) = 0.07, *p* = 0.608) for faces.

We then analyzed whether novel and best alternative choices are associated with distinct constructs of a participant’s social network. We calculated the proportion of individual probabilities for novel and best choices, with higher values in this score reflecting higher undirected novelty seeking when compared with choosing faces with known rewards. The more often a participant chose novel face pictures instead of familiar ones, the larger the social network size of the participant, as indexed by the Lubben Social Network index (**Figure 2D**, Partial R(19) = 0.67, *p* = 0.001) and the Social Network Index (**Figure 2E**, Partial R(19) = 0.64, *p* = 0.002). In contrast, when investigating indicators of social network quality, the ratio of novel versus familiar choices did not correlate with social connectedness (Social Connectedness Scale, Partial R(19) = 0.24, *p* = 0.301) nor with perceived loneliness (UCLA-20, Partial R(19) = −0.26, *p* = 0.253).

### III.c. Novel over familiar face selection activates the supramarginal and mid-cingulate cortices

Participant’s trial-by-trial choices, within two trials from an insertion, were used as regressors in the fMRI analysis to identify brain activity related to novel and best alternative choices (**Figure 2F**, height threshold *p* < 0.005 and a cluster-level FDR-corrected threshold of *p* < 0.05, **Table 2**). The contrast novel > best revealed increased activation in a distributed set of regions encompassing the left and right supramarginal cortices, the mid-cingulate, and primary sensorimotor and supplementary motor cortices. No clusters survived the chosen significance threshold for the novel < best contrast.

**Table 2.** Result table of fMRI findings. Cluster size, *t*-value, and coordinates of peak voxel, with associated brain region based on the Harvard-Oxford probabilistic atlas.

| Figure 1F - faces > fixation cross |  |  |  |
| --- | --- | --- | --- |
| Cluster size (voxels) | t-value | Coordinates (x, y, z) | Harvard-Oxford label |
| 69771 | 8.83 | (-32, -44, -18) | Left Temporal Occipital Fusiform Cortex |
| 562 | 6.14 | (30, 22, 2) | Right Insular Cortex |
| 64 | 5.36 | (32, 62, -14) | Right Frontal Pole |
| 1257 | 5.32 | (6, 30, 24) | Right Cingulate Gyrus anterior division |
| 43 | 5.19 | (-8, -14, -40) | Brainstem |
| 48 | 5 | (-34, -8, -38) | Left Temporal Fusiform Cortex anterior division |
| 138 | 4.93 | (20, 18, -32) | Unspecified |
| 426 | 4.81 | (-38, 50, -2) | Left Frontal Pole |
| 56 | 4.64 | (12, 30, -22) | Right Frontal Orbital Cortex |
| 66 | 4.62 | (6, -16, 30) | Right Cingulate Gyrus posterior division |
| 74 | 4.27 | (32, -2, -38) | Right Temporal Fusiform Cortex anterior division |
| 53 | 4.01 | (-8, 4, 2) | Left Caudate |
| 48 | 3.95 | (34, 50, 2) | Right Frontal Pole |
| 32 | 3.7 | (34, 62, -2) | Right Frontal Pole |
| Figure 1G - reward > no reward |  |  |  |
| Cluster size (voxels) | t-value | Coordinates (x, y, z) | Harvard-Oxford label |
| 2666 | 8.12 | (-14, -46, 36) | Left Cingulate Gyrus posterior division |
| 1967 | 8.04 | (0, 50, 0) | Left Paracingulate Gyrus |
| 1214 | 6.43 | (-22, 8, -10) | Left Putamen |
| 777 | 6.05 | (-12, -100, 22) | Left Occipital Pole |
| 1344 | 5.76 | (50, -26, 18) | Right Parietal Operculum Cortex |
| 1943 | 5.58 | (-60, -28, 24) | Left Supramarginal Gyrus anterior division |
| 92 | 5.43 | (-56, 10, -18) | Left Temporal Pole |
| 434 | 5.31 | (-24, 28, -14) | Left Frontal Orbital Cortex |
| 320 | 5.28 | (-64, -10, -10) | Left Middle Temporal Gyrus anterior division |
| 90 | 5.11 | (44, 2, 10) | Right Central Opercular Cortex |
| 41 | 4.96 | (22, -34, 48) | Unspecified |
| 310 | 4.94 | (42, -24, 58) | Right Postcentral Gyrus |

| 71 | 4.9 | (-10, -28, 18) | Left Lateral Ventricle |
| --- | --- | --- | --- |
| 244 | 4.88 | (16, -102, 10) | Right Occipital Pole |
| 133 | 4.64 | (12, -46, 62) | Right Postcentral Gyrus |
| 174 | 4.55 | (-46, -82, 2) | Left Lateral Occipital Cortex inferior division |
| 529 | 4.36 | (-14, 34, 52) | Left Superior Frontal Gyrus |
| 30 | 4.33 | (14, 44, 58) | Unspecified |
| 94 | 4.27 | (-52, 12, 12) | Left Inferior Frontal Gyrus pars opercularis |
| 30 | 4.01 | (20, 14, 14) | Right Caudate |
| 25 | 3.87 | (38, -6, -16) | Right Insular Cortex |
| 55 | 3.85 | (-14, -82, 42) | Left Lateral Occipital Cortex superior division |
| 134 | 3.82 | (-30, -24, -14) | Left Hippocampus |
| 25 | 3.79 | (28, -16, -18) | Right Hippocampus |
| 31 | 3.75 | (-48, 20, -32) | Left Temporal Pole |
| 39 | 3.63 | (2, -50, -42) | Unspecified |
| 34 | 3.48 | (10, -92, 38) | Right Occipital Pole |
| 70 | 3.34 | (24, -80, -40) | Unspecified |
| 25 | 3.34 | (-4, 60, 38) | Left Frontal Pole |
| Figure 2F - novel > best |  |  |  |
| Cluster size (voxels) | t-value | Coordinates (x, y, z) | Harvard-Oxford label |
| 263 | 5.56 | (50, -46, -6) | Right Inferior Temporal Gyrus temporooccipital part |
| 577 | 5.45 | (-52, -24, 32) | Left Supramarginal Gyrus anterior division |
| 2650 | 5.39 | (-2, -32, 50) | Left Precentral Gyrus |
| 31 | 5.15 | (26, -24, 12) | Unspecified |
| 61 | 4.87 | (62, 14, 22) | Right Precentral Gyrus |
| 358 | 4.84 | (-48, 0, 6) | Left Central Opercular Cortex |
| 269 | 4.84 | (40, -10, 58) | Right Precentral Gyrus |
| 103 | 4.74 | (34, -54, 64) | Right Superior Parietal Lobule |
| 112 | 4.47 | (18, -34, 62) | Right Postcentral Gyrus |
| 107 | 4.37 | (36, -24, -28) | Right Temporal Fusiform Cortex posterior division |
| 64 | 4.31 | (40, -58, -20) | Right Temporal Occipital Fusiform Cortex |
| 44 | 4.21 | (68, -6, -18) | Right Middle Temporal Gyrus |

|  |  |  | anterior division |
| --- | --- | --- | --- |
| 212 | 4.19 | (62, -16, 24) | Right Postcentral Gyrus |
| 76 | 3.94 | (34, -12, 8) | Right Insular Cortex |
| 35 | 3.91 | (32, -42, 72) | Right Superior Parietal Lobule |
| 29 | 3.9 | (34, -78, -2) | Right Lateral Occipital Cortex<br>inferior division |
| 29 | 3.87 | (10, 2, 40) | Right Cingulate Gyrus anterior<br>division |
| 44 | 3.85 | (-52, -62, 16) | Left Lateral Occipital Cortex<br>superior division |
| 60 | 3.76 | (46, -20, 62) | Right Postcentral Gyrus |
| 35 | 3.76 | (-16, -22, 10) | Left Thalamus |
| 26 | 3.66 | (30, -42, -40) | Unspecified |
| 58 | 3.63 | (-6, -32, 38) | Left Cingulate Gyrus posterior<br>division |
| 43 | 3.6 | (52, -78, 12) | Right Lateral Occipital Cortex<br>inferior division |
| 48 | 3.6 | (68, -40, -12) | Right Middle Temporal Gyrus<br>temporooccipital part |
| 31 | 3.57 | (38, -84, 28) | Right Lateral Occipital Cortex<br>superior division |
| 29 | 3.32 | (62, -46, -14) | Right Inferior Temporal Gyrus<br>temporooccipital part |
| Figure 2G - reward > no reward |  |  |  |
| Cluster size (voxels) | t-value | Coordinates (x, y, z) | Harvard-Oxford label |
| 419 | 6.76 | (34, -4, -22) | Right Amygdala |
| 78 | 4.89 | (-66, -2, -4) | Unspecified |
| 172 | 4.8 | (34, -22, 56) | Right Precentral Gyrus |
| 79 | 4.7 | (-14, -46, 38) | Left Precuneus Cortex |
| 63 | 4.43 | (2, -20, 70) | Right Precentral Gyrus |
| 60 | 4.39 | (26, 8, -10) | Right Putamen |
| 72 | 4.34 | (-4, -32, 38) | Left Cingulate Gyrus posterior<br>division |
| 93 | 4.24 | (4, -50, -42) | Unspecified |
| 35 | 4.15 | (-30, -6, -22) | Left Amygdala |
| 129 | 3.98 | (-24, -90, 12) | Left Occipital Pole |
| 27 | 3.95 | (-12, -26, 72) | Left Precentral Gyrus |
| 42 | 3.94 | (2, 22, -10) | Right Subcallosal Cortex |

| 79 | 3.87 | (-8, -40, 72) | Left Postcentral Gyrus |
| --- | --- | --- | --- |
| 49 | 3.81 | (44, -30, 18) | Right Parietal Operculum Cortex |
| 46 | 3.77 | (22, -34, 72) | Right Postcentral Gyrus |
| 28 | 3.68 | (16, 44, 60) | Unspecified |
| 29 | 3.65 | (-2, -52, 16) | Left Cingulate Gyrus posterior division |
| 45 | 3.51 | (-22, -90, -6) | Left Occipital Fusiform Gyrus |
| 27 | 3.4 | (14, -24, 72) | Right Precentral Gyrus |
| Figure 3F - explore > exploit |  |  |  |
| Cluster size (voxels) | t-value | Coordinates (x, y, z) | Harvard-Oxford label |
| 157 | 4.4 | (-6, -56, 40) | Left Precuneus Cortex |
| 50 | 4.17 | (-8, 22, 68) | Left Superior Frontal Gyrus |
| 70 | 3.94 | (36, -6, 60) | Right Precentral Gyrus |
| 147 | 3.85 | (-6, 26, 34) | Left Paracingulate Gyrus |
| 40 | 3.7 | (-10, 40, 48) | Left Frontal Pole |
| 48 | 3.55 | (46, -68, 18) | Right Lateral Occipital Cortex superior division |
| 65 | 3.54 | (-48, 10, 40) | Left Middle Frontal Gyrus |
| 36 | 3.45 | (52, -42, -24) | Right Inferior Temporal Gyrus temporooccipital part |
| 63 | 3.42 | (-50, 20, 18) | Left Inferior Frontal Gyrus pars opercularis |
| 40 | 3.38 | (42, -84, 20) | Right Lateral Occipital Cortex superior division |
| Figure 3G - RPE pos > RPE neg |  |  |  |
| Cluster size (voxels) | t-value | Coordinates (x, y, z) | Harvard-Oxford label |
| 151 | 5.71 | (36, 64, 8) | Right Frontal Pole |
| 105 | 5.39 | (-52, -42, 62) | Unspecified |
| 214 | 5.2 | (-38, 34, 34) | Left Middle Frontal Gyrus |
| 85 | 4.9 | (22, 50, 16) | Right Frontal Pole |
| 141 | 4.66 | (-34, -70, 26) | Left Lateral Occipital Cortex superior division |
| 246 | 4.63 | (50, 34, 32) | Right Middle Frontal Gyrus |
| 66 | 4.54 | (-34, -54, 70) | Unspecified |
| 424 | 4.46 | (-34, 66, 8) | Left Frontal Pole |
| 26 | 4.46 | (-24, 14, 30) | Unspecified |
| 46 | 4.01 | (-26, 34, -2) | Unspecified |
| 42 | 3.92 | (40, -64, 32) | Right Lateral Occipital Cortex superior division |
| 28 | 3.91 | (38, 28, 54) | Right Middle Frontal Gyrus |
| 63 | 3.72 | (16, -76, 58) | Right Lateral Occipital Cortex<br>superior division |
| 63 | 3.67 | (58, -50, -10) | Right Middle Temporal Gyrus<br>temporooccipital part |
| 91 | 3.55 | (-14, -76, 58) | Left Lateral Occipital Cortex<br>superior division |
| 37 | 3.36 | (32, -86, 42) | Right Lateral Occipital Cortex<br>superior division |
| 44 | 3.34 | (-2, -62, 58) | Left Precuneus Cortex |
| 30 | 3.33 | (-24, 50, -8) | Left Frontal Pole |

We further explored brain activity changes associated with rewards and no-rewards for choices made within two trials from a novel insertion (**Figure 2G**, height threshold *p* < 0.005 and a cluster-level FDR-corrected threshold of *p* < 0.05, **Table 2**). The reward > no-reward contrast revealed activations in bilateral anterior hippocampus, the precuneus, the right ventral putamen, and the basal forebrain. The reward < no-reward contrast revealed a unique small cluster of deactivations in the left superior frontal cortex.

### III.d. Face exploration correlates with social connectedness

Trial-by-trial estimates for the IEV and the exploration BONUS of a choice were derived from the POMDP model. IEV was higher for best alternative choices when compared to novel choices, with the IEV of novel choices increasing as the number of trials since a novel insertion increased (**Figure 3A**). BONUS was higher for novel choices when compared to best alternative choices, with the BONUS of novel choices decreasing as the number of trials since a novel insertion increased (**Figure 3B**). Plotting trial-by-trial BONUS versus IEV estimates was used to identity exploratory choices (BONUS>0 & IEV<0.5), exploitative choices (BONUS<0 & IEV>0.5) as well as choices that were neither exploratory nor exploitative (**Figure 3C**). Aligned with previous studies, exploration peaked when novel choices were inserted and quickly decreased as the number of trials since a novel insertion increased, with exploitation increasing instead (**Figure 3D**). Averaged probabilities of exploration, exploitation, and neither choices were calculated for each participant, revealing that overall participants engaged more often in exploitation during the duration of the whole task, with exploration being the least selected choice (**Figure 3E**, F(2,48) = 3.44, *p* < 0.05, post-hoc test *p*<0.05 uncorrected and FWE corrected). Individual choice probabilities for exploration, exploitation, and neither were correlated with individual probabilities of novel, best, and worst choices within two trials from an insertion, showing strong internal consistency across best-exploitation, novel-exploration, and worst-neither choices (**Figure 3F**).

**Figure 3.**
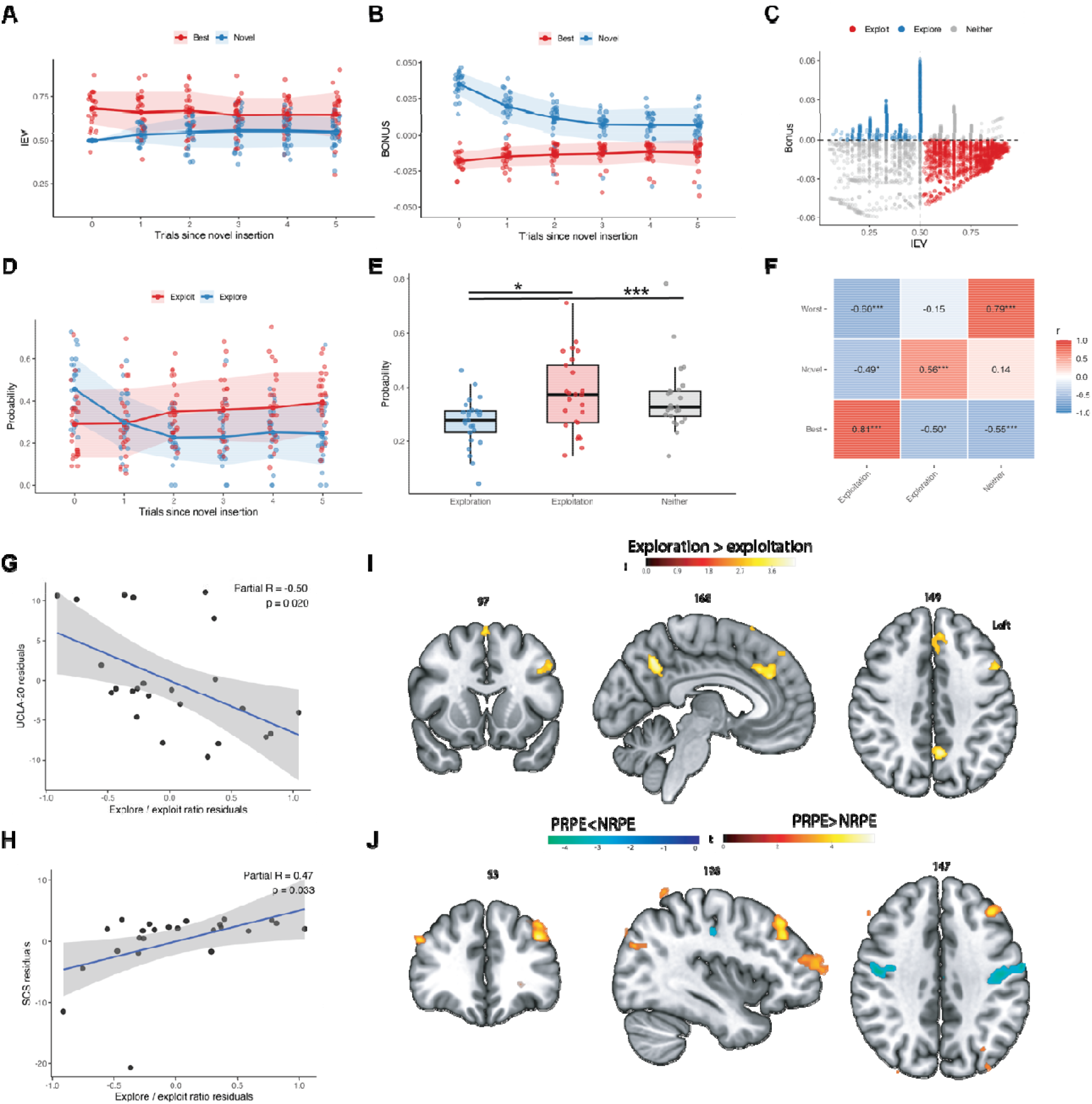
Face exploration predicts social network quality and is associated with activity in anterior cingulate, precuneus, and lateral prefrontal cortex. (A-B) Mean trial-by-trial changes in POMDP estimates of IEV and BONUS during the selection of novel options (in blue) versus the best alternative choice (in red). (C) Trials were classified as explorative, exploitative, or neither depending on their corresponding IEV and BONUS values. (D) Participants explored less often as the number of trials elapsed since a new insertion was introduced, instead, increasing exploitative behavior. (E) Over the whole task, participants engaged more frequently in exploitation, with exploration being the least frequent behavior. (F) Modelled exploitative, explorative, and unclassified behavior (neither) correlated positively with the individual probabilities of choosing best, novel, and worst options, respectively, suggesting consistency across the modelled and empirical behaviors. **(G-H)** The ratio of explorative versus exploitative behavior correlated with two measures of social network quality, namely negatively with loneliness as assessed through the UCLA-20, and positively with social connectedness as assessed through the Social Connectedness Scale (SCS). Partial Pearson’s correlation coefficients corrected for age, gender, and years of education. **(I)** Increased activity in the anterior cingulate cortex and precuneus (warm colors) for exploration versus exploitation. **(J)** The positive reward prediction error (PRPE) versus the negative reward prediction error (NRPE) revealed activation in the bilateral prefrontal cortices (hot colors), while deactivations were identified in the somatosensory cortex (cold colors). Significant clusters were identified with a height threshold *p* < 0.005 and a cluster-level FDR-corrected threshold of *p* < 0.05; color bars show *t*-values. BONUS = exploration bonus; IEV = immediate expected value; POMDP = partially observable Markov decision making process. \**p* < 0.05; ***FWE-corrected *p* < 0.05

We next analyzed whether explorative and exploitative choices were associated with distinct constructs of a participant’s social network. We calculated the proportion of individual probabilities for explorative and exploitative choices, with higher values in this score reflecting higher goal-directed exploration when compared to exploitation. The more often a participant explored new faces instead of exploiting faces with known rewards, the less lonely participants felt as indexed by the UCLA-20 (**Figure 3G**, Partial R(19) = −0.50, *p* = 0.020), and the more socially connected they felt as indexed by the Social Connectedness Scale (**Figure 3H**, Partial R(19) = 0.47, *p* = 0.033). In contrast to the novelty/familiar choices findings, the ratio of exploration versus exploitation did not correlate with measures of social network size (Social Network Index, Partial R(19) = 0.38, *p* = 0.090; Lubben Social Network index, Partial R(19) = 0.35, *p* = 0.125).

### III.e. Face exploration over exploitation activates the anterior cingulate and the left dorsolateral prefrontal cortex

Participant’s trial-by-trial choices derived from the POMDP model were used as regressors in the fMRI analyses to identify brain activity related to explorative and exploitative choices (**Figure 3I**, height threshold *p* < 0.005 and a cluster-level FDR-corrected threshold of *p* < 0.05, **Table 2**). The contrast exploration > exploitation revealed increased activation in a distributed set of regions encompassing anterior cingulate, precuneus, and left dorsolateral prefrontal cortex. No clusters survived the chosen significance threshold for the exploration < exploitation contrast.

We further explored brain activity changes associated with PRPE and NRPE (**Figure 3J**, height threshold *p* < 0.005 and a cluster-level FDR-corrected threshold of *p* < 0.05, **Table 2**). The PRPE > NRPE contrast revealed bilateral activations in the lateral prefrontal cortices and the precuneus, whereas the PRPE < NRPE contrast revealed a larger bilateral cluster of deactivations in the somatosensory cortices.

### III.f. Social and decision-making circuits

To further characterize the functional profile of brain activity associated with explorative and novel choices, we examined whether neural differentiation between choice types was preferentially expressed within brain systems associated with cognitive processes typically involved in a multi-arm bandit task. Specifically, we compared the magnitude of subject-level absolute activity differences across independently defined Neurosynth maps linked to the terms “social”, “decision making”, and “working memory” (**Figure 4A-C**). Neural differentiation varied significantly across the three cognitive networks for both exploration versus exploitation and novel versus best choices (*F*(3,72) = 19.75, *p* < 0.001; **Figure 4D-E**). Post-hoc paired comparisons indicated greater differentiation within social and decision-making networks relative to working memory for both the exploration versus exploitation and novel versus best choices comparisons (repeated-measures *p* < 0.05 Holm-corrected; **Figure 4D-E**).

**Figure 4.**
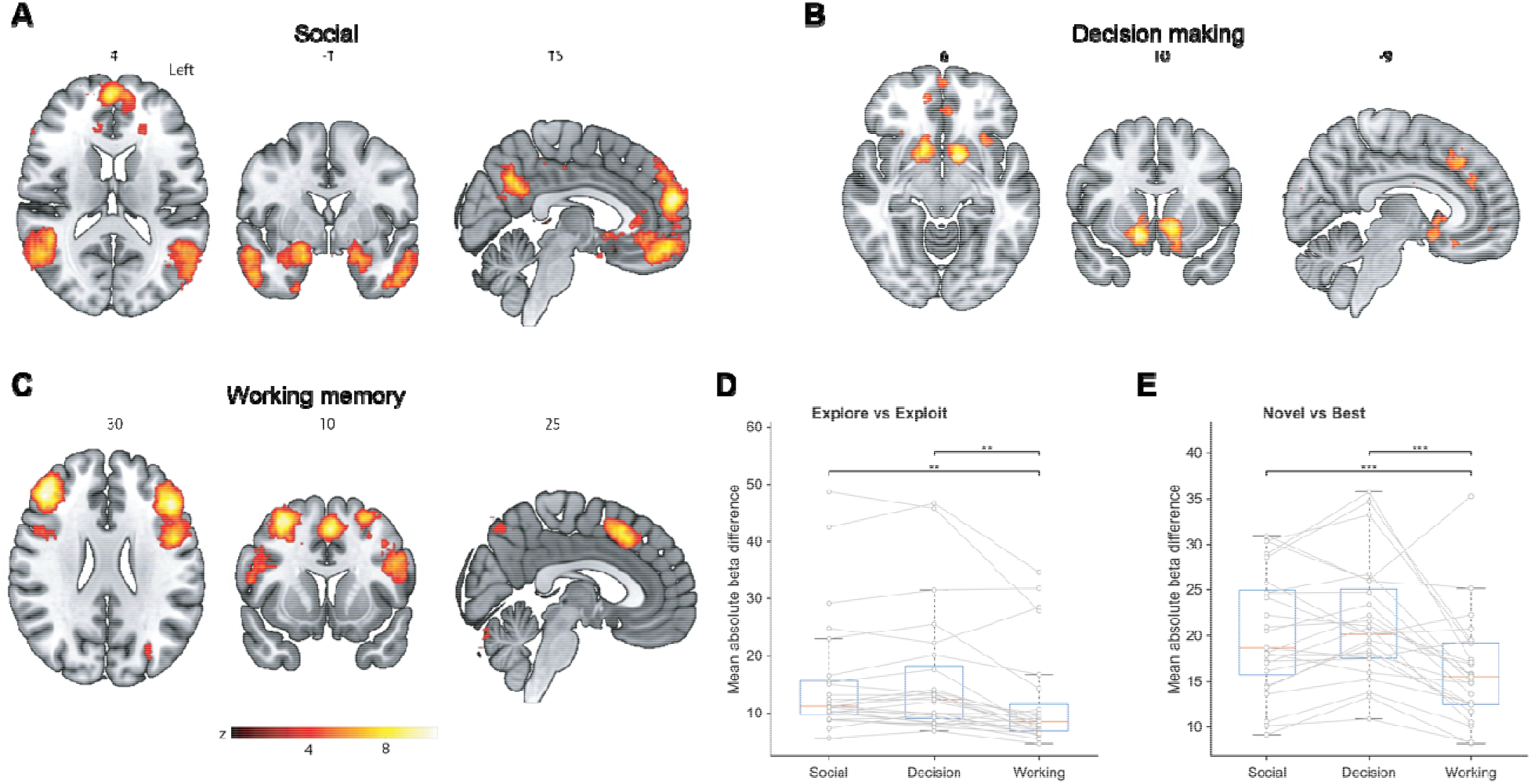
Exploration- and novelty-related circuit differentiation within Neurosynth-defined cognitive networks. (A–C) FDR-thresholded Neurosynth association maps for social cognition, decision making, and working memory, respectively (FDR corrected *p* < 0.01); warm colors indicate higher Neurosynth *z*-values. **(D)** Mean absolute differences in brain activity between exploration and exploitation (left) and between novel and best choices (right), calculated within each Neurosynth-defined network for each participant. Boxplots show the median and interquartile range, with individual participant values connected across networks. A larger mean absolute beta difference indicates greater neural differentiation between the corresponding behavioral conditions, irrespective of the direction of the effect. *p* < 0.05; \**p* < 0.01; \*\**p* < 0.001, Holm-corrected.

### III.g. Non-social decision making does not correlate with social network constructs

To assess whether behavioral effects observed during the social three-arm bandit task generalized to a non-social context, 16 participants completed a remote version of the task in which human faces were replaced by pictures of animals. As in the social version of the task, participants became more likely to choose best options as the number of trials following the introduction of a novel option increased. However, the probability of choosing the novel option remained stable across trials (**Figure 5A**). During the first two trials following a novel insertion, participants selected the best option more frequently than the novel option, with the worst option selected least frequently (**Figure 5B**, *F(2,30) = 9.01, p < 0.005)*. In contrast to the social version of the task, the novel/best choice ratio in the non-social task was not significantly associated with either measure of social network size (SNI or LSN; **Figure 5C**). Correlations with social network size remained significant in this subsample for the social version of the task, although the strength of these correlations did not differ significantly between the social and non-social tasks (**Figure 5C**).

**Figure 5.**
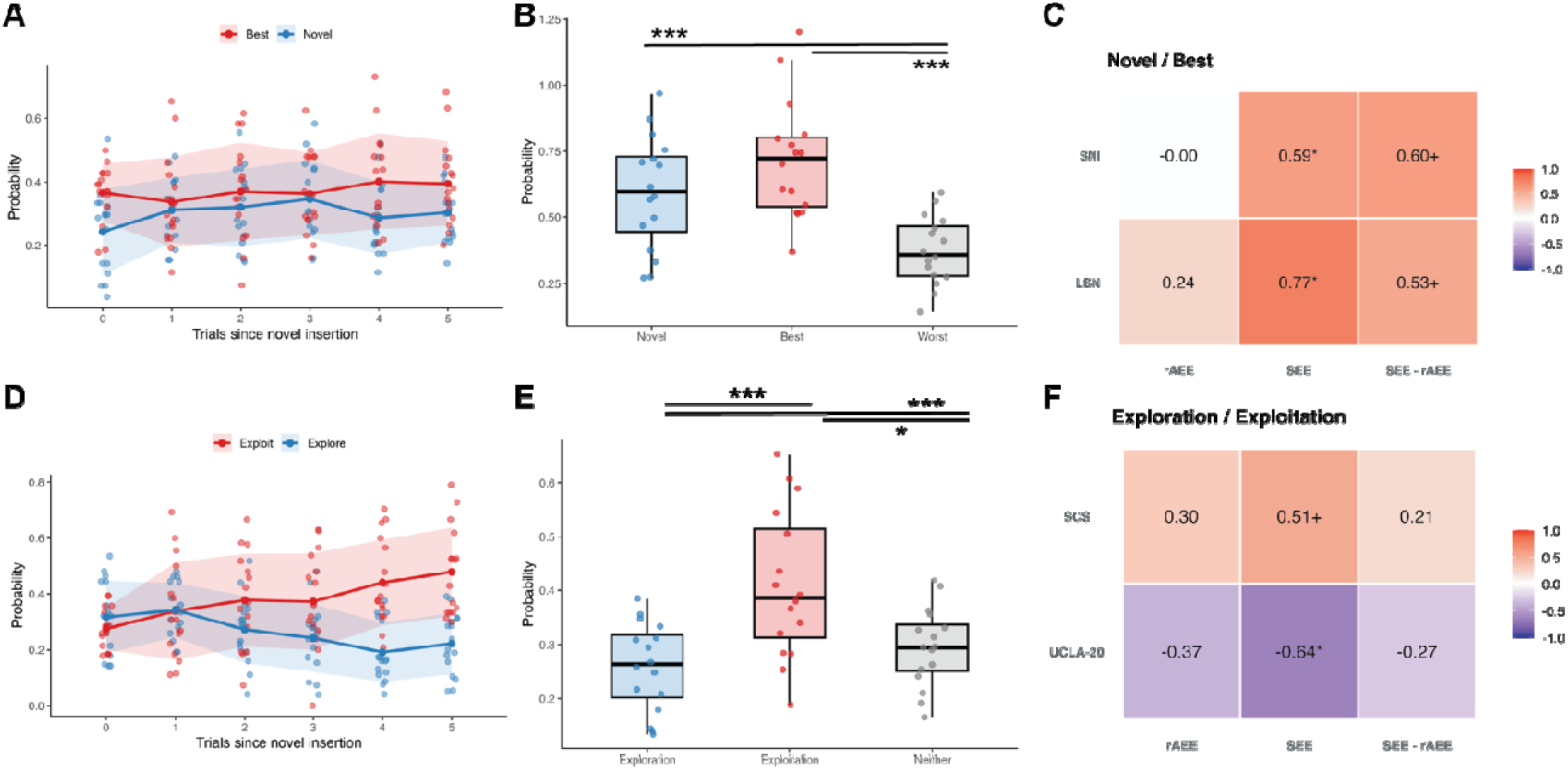
Non-social version of the three-arm bandit-task. 16 participants completed a non-social version of the three-arm bandit-task remotely, which showed pictures of animals as stimuli instead of human faces. **(A)** Participants choose best options more often as the number of trials elapsed since a new insertion increased, while the likelihood of choosing novel options remained relatively stable. **(B)** The probabilities of choosing the novel, best, and worst option were computed for the first two trials after a novel insertion. In contrast to the social version of the task, best options were selected more often than novel options, with worst options being selected the least frequently. **(C)** In this subsample, **t**he ratio of novel versus best options did not significantly correlate with two measures of social network size (LSN and SNI; rAEE, left column), while significant correlations were retained for the social version of the task (SEE, middle column). The difference in correlations between both tasks was not significant (SEE-rAEE, right column). **(D)** Participants explored less often as the number of trials elapsed since a new insertion was introduced, instead, increasing exploitative behavior. **(E)** Similarly to the social version of the task, over the whole task participants engaged more frequently in exploitation, with exploration being the least frequent behavior. **(F)** In this sample, the exploration/exploitation ratio did not correlate significantly with loneliness (UCLA-20) or social connectedness (SCS) in the non-social version of the task (rAEE, left column), although it retained significant and trending correlations for the social version of the bandit-task (SEE, middle column). The difference in correlations between both tasks was not significant (SEE-rAEE, right column). rAEE = remote non-social version of the three-arm bandit task showing animal pictures; SEE = social version of the three-arm bandit task showing pictures of older adults. ^+^*p* < 0.1; \**p* < 0.05; ***FWE-corrected *p* < 0.05

Across the non-social task, exploration similarly decreased and exploitation increased with increasing distance from the introduction of a novel option (**Figure 5D**), with participants overall engaging more frequently in exploitation than exploration (**Figure 5E**, *F(2,30) = 6.56, p < 0.005*). The exploration/exploitation ratio was not significantly associated with loneliness (UCLA-20) or social connectedness (SCS) in the non-social task (**Figure 5F**). In contrast, the corresponding associations remained significant or trend-level in this subsample for the social task, although direct comparisons indicated no significant differences in correlation strength between the social and non-social tasks (**Figure 5F**).

Choice probabilities for best, novel, exploration, and exploitation in the non-social version of the task did not significantly correlate with corresponding measures in the social task version (Partial R_best_ (10) = −0.04, p = 0.91; Partial R_novel_ (10) = 0.17, p = 0.59; Partial R_exploitation_ (10) = 0.37, p = 0.28; Partial R_exploration_ (10) = 0.29, p = 0.35).

## IV. Discussion

In this social decision-making neuroimaging study in older adults, we identified dissociable social constructs and brain signatures of undirected novelty seeking versus goal-directed exploration. While older adults choosing novel inserted faces over faces with known rewards had larger social networks and increased activity in posterior brain areas, goal-directed exploration was associated with higher social connectedness and activated anterior brain areas. This study elucidates how older adults deploy complementary social decision-making strategies when facing options with uncertain outcomes, and how these choices relate to: *(i)* distinct constructs of social functioning and *(ii)* rely on different neural systems. These findings have important implications, suggesting that different strategies may be needed to target and prevent social isolation and loneliness in older adults, two major risk factors contributing to worsening mental and physical health in aging (Cacioppo et al., 2010; Donovan & Blazer, 2020).

### IV.a. Decision-making strategies and social constructs in older adults

A common challenge in everyday life is balancing the pursuit of novel opportunities against familiar and reliable options (Cohen et al., 2007; Hills et al., 2015). This dilemma, known as the exploration/exploitation tradeoff, requires weighting options with well-known reward values against options with an uncertain but potentially higher payoff (Averbeck, 2015). Age-related exploitation biases in older adults have been proposed to result from the accumulation of prior knowledge and from a need to optimize affective goals (Carstensen, 2021; Spreng & Turner, 2021). Consistent with this view, older adults often prioritize smaller, emotionally closer social networks to support their emotional well-being, when compared with younger adults (Carstensen, 1992; English & Carstensen, 2014). Previous work suggests that the pursuit of novel opportunities is governed by two separable processes: goal-directed exploration, which prioritizes information acquisition in the face of uncertainty, and random undirected novelty seeking, which reflects stochastic fluctuations in choice when presented with an unseen option (Cockburn et al., 2022; Wilson et al., 2014). Although overlapping, undirected novelty seeking and goal-directed exploration reflect distinct decision-making strategies, either reflecting the sudden attendance to novel, salient stimuli versus having to strategize between options with uncertain outcomes (Hogeveen et al., 2022). A crucial question is whether individual tendencies toward exploitation, undirected novelty seeking, and goal-directed exploration reflect stable trait-like characteristics or more transient state-dependent processes. If these decision-making strategies represent stable traits, they may constitute behavioral phenotypes that shape resilience and vulnerability across the lifespan, representing promising targets for behavioral, social, or pharmacological interventions. Longitudinal studies will be necessary to determine the stability of these behaviors over time and to assess their potential as markers of healthy and pathological aging.

In this context, the specific association of novelty seeking with larger social networks and of exploration with increased connectedness may have important implications for understanding social and affective functioning in later life. One possible interpretation is that undirected novelty seeking facilitates the acquisition of new social contacts, thereby influencing network size, whereas goal-directed exploration may support the evaluation and maintenance of social relationships, thereby influencing subjective connectedness. Although often co-occurring, social isolation is defined as the objective lack of social connections (Holt-Lunstad, 2024), differing from loneliness, defined as the subjective, distressing feeling of being alone or disconnected (Cacioppo et al., 2014). Both constructs have been shown to contribute to adverse, although dissociable, health outcomes (Coyle & Dugan, 2012; Holt-Lunstad et al., 2015). Loneliness is more strongly associated with deleterious mental health outcomes, including depression and reduced meaning in life (Ge et al., 2017), whereas objective isolation is a stronger predictor of general physical decline and early mortality (Holt-Lunstad et al., 2015; Steptoe et al., 2013). These observations raise the possibility that novelty seeking and social exploration may contribute to distinct dimensions of social functioning in later life. If replicated, interventions that promote engagement with novel social experiences may preferentially influence objective social isolation, whereas interventions targeting the valuation of social opportunities may more strongly affect loneliness and perceived connectedness. Consistent with this possibility, a meta-analysis of loneliness interventions found that modifying how individuals evaluate social opportunities was more effective at reducing loneliness than simply increasing social contact (Masi et al., 2011). Future studies directly manipulating undirected novelty seeking behavior and goal-directed social exploration in older adults will be necessary to test these hypotheses and establish causal relationships.

### IV.b. Dissociable neural systems for social exploration and novelty seeking

A growing body of research has investigated the neural mechanisms underlying age-related shifts in explore–exploit behavior, proposing that age-related changes in medial temporal, medial parietal, and prefrontal systems supporting memory, prospection, and value-based decision-making contribute to a bias toward exploitation at the expense of exploration in older adults (Spreng & Turner, 2021). In parallel, distinct brain systems have been shown to underlie the detection of behaviorally relevant stimuli, particularly when they are salient or unexpected, versus the goal-directed, top-down modulated selection of stimuli and responses (Corbetta & Shulman, 2002). In the non-social context, temporoparietal, inferior frontal, and dopaminergic midbrain–striatal circuits have been repeatedly involved in salience detection and orienting attention towards novel stimuli. Aligned with these previous reports, our study revealed that choosing novel options versus the best rewarding option activates a set of posterior brain regions including the mid-cingulate cortex, supramarginal gyrus, and supplementary motor areas. The supramarginal activations found in our analyses are consistent with previous studies reporting temporoparietal activations to novel faces, a finding believed to reflect the need to interpret social cues, evaluate emotional expressions, and build social concepts (Molenberghs et al., 2016; Schurz et al., 2014). Rewards associated with these early choices engaged a striatal circuit, in line with models implicating dopaminergic signaling in undirected novelty seeking behavior (Bunzeck & Düzel, 2006; Wittmann et al., 2008). Together, these findings suggest that posterior salience-related and striatal reward circuits may support social novelty seeking by facilitating the detection and pursuit of new social opportunities.

Conversely, goal-directed exploration has been shown to activate a distributed brain system comprising the dorsal anterior cingulate, as well as the medial and lateral prefrontal cortex (Daw et al., 2006; Hogeveen et al., 2022). Our findings are well aligned with these previous reports, as evidenced by increased activity in the anterior cingulate cortex and lateral prefrontal cortices in social exploration when compared to social exploitation. When comparing positive reward prediction errors with negative reward prediction errors, our analyses revealed increased activity in the bilateral prefrontal cortex, a set of regions playing a crucial role in the updating of internal beliefs and in supporting adaptive decision-making processes of goal-directed exploration (Badre et al., 2012; Gläscher et al., 2010). Previous research has demonstrated a causal role for the dorsolateral prefrontal cortex in belief updating under uncertainty (Schulreich & Schwabe, 2021), both in social and non-social contexts, enabling decision-making behavior to flexibly adjust following unexpected outcomes. Complementary evidence has shown that dmPFC activity is enhanced during social interactions requiring representations of others’ mental states (Amodio & Frith, 2006; Christian et al., 2024). Taken together, these findings support the view that social exploration relies on anterior control networks involved in belief updating and adaptive decision-making, complementing the posterior salience- and reward-related systems that support social novelty seeking.

### IV.c. Limitations and future directions

A limitation of the study is the use of faces as only social stimuli. As such, we cannot exclude the possibility that non-social features of the faces used as stimuli contributed to choice selections. This concern is partially mitigated by the following factors: *(i)* choice selection did not correlate with out-of-sample scores of the face’s likeability and trustworthiness; *(ii)* probabilities of reward were randomly attributed to faces for each participant; and *(iii)* choice ratios from a non-social version of the task did not correlate with choices from the social task nor with social constructs. Future studies could integrate non-social and more diverse social stimuli (e.g., gatherings, interaction between persons) (Isik et al., 2017) as well as use more socially salient reward cues (Jones et al., 2011; Ruff & Fehr, 2014) or manipulations of social beliefs (Behrens et al., 2008; Hackel et al., 2015). These study design choices could be implemented in larger, better-powered studies spanning multiple age groups (Blanco & Sloutsky, 2024; Somerville et al., 2017; Spreng & Turner, 2019).

Furthermore, the study of social-affective decision making and its neural correlates would likely benefit from the inclusion of real-word indicators of social behavior. For example, ecological momentary assessments and global positioning data acting as proxies of regular social interactions (Choi et al., 2025; Fingerman et al., 2020) and social mobility (Crane et al., 2023; Yu et al., 2025), could contribute to the development of ecologically valid phenotypes of social decision-making in later life. Future studies could extend the investigation of changes in social decision-making strategies and their neural correlates to clinical populations, including late-life depression and dementia (Holt-Lunstad, 2024). Such studies could contribute to the identification of behavioral and neural phenotypes of social-decision making and the development of prevention-oriented approaches supporting adaptive social engagement for healthy aging.

## Data availability statement

Analyses were performed in MATLAB 2023b, Python, and R. Code will be available on the following GitHub page (https://cvaltierra-neuro.github.io/) in addition to de-identified behavioral and secondary imaging data upon acceptance of the manuscript.

## Author contributions

LP and AG conceived the study. GM, SM, AO, SG, CA, and CV collected the data and performed the experiments. NS and GT contributed tools to curate and collect data. CV, JC, PMC, and LP analyzed the data. JH supervised data analysis and provided tools to analyze the data. JM provided critical feedback for the interpretation of the findings. LP and CV wrote the original draft of the manuscript. All authors read, edited, and approved the manuscript.

## Acknowledgements

We thank the participants of the study for their invaluable contribution to research, and Amy Markowitz for her constructive feedback on the manuscript.

